# Emergence of temporal noise hierarchy in co-regulated genes of multi-output feed-forward loop

**DOI:** 10.1101/2024.08.08.607134

**Authors:** Mintu Nandi

## Abstract

Natural variations in gene expression, called noise, are fundamental to biological systems. The expression noise can be beneficial or detrimental to cellular functions. While the impact of noise on individual genes is well-established, our understanding of how noise behaves when multiple genes are co-expressed by shared regulatory elements within transcription networks remains elusive. This lack of understanding extends to how the architecture and regulatory features of these networks influence noise. To address this gap, we study the multi-output feed-forward loop motif. The motif is prevalent in bacteria and yeast and influences co-expression of multiple genes by shared transcription factors. Focusing on a two-output variant of the motif, the present study explores the interplay between its architecture, co-expression patterns of the two genes (including symmetric and asymmetric expressions), and the associated noise dynamics. We employ a stochastic modeling approach to investigate how the binding affinities of the transcription factors influence symmetric and asymmetric expression patterns and the resulting noise dynamics in the co-expressed genes. This knowledge could guide the development of strategies for manipulating gene expression patterns through targeted modulation of transcription factor binding affinities.

## I. INTRODUCTION

Genes are the blueprints of proteins. The information available in a gene gets translated to protein through gene expression. Gene expression involves two steps: transcription of the gene into mRNA and translation of the resulting mRNA into protein [1–4]. Gene expression is tightly controlled within cells to produce the right amounts of proteins in a timely manner. Precise regulation and accuracy of gene expression are necessary for cell fitness and development [1]. Such control of gene expression happens throughout various stages of the expression process. One key mechanism to control precise gene expression is transcriptional regulation, where a specific transcription factor (TF), a protein, binds to a particular site in the promoter region [5–8]. The TF can act as an activator, recruiting the machinery (e.g., RNA polymerase) needed for transcription and promoting the gene expression rate. On the other hand, it can be a repressor that blocks the transcriptional machinery, thus impeding gene expression [9]. The role of TFs in transcriptional regulation orchestrates gene transcription regulatory networks (GTRNs) where multiple patterns of interaction among the TFs and genes emerge [10]. In GTRNs, nodes represent TFs and genes, and edges denote the regulation of the genes by TFs [9,10].

Gene expression noise is inherent and ubiquitous in output proteins due to the stochastic nature of the reaction steps involved [11]. The quantification of noise and identification of its various sources in simple gene expression and GTRNs are well-established [6,8,11–17]. Feed-forward loop (FFL) is such a recurring regulatory motif, frequently occurs in GTRN of *Escherichia coli* and *Saccharomyces cerevisiae* [18–21]. The coherent FFLs, a specific class of FFLs, are prevalent in *E. coli*, yeast, the drosophila nervous system, and other higher organisms [9,22–24]. Numerous experimental studies have investigated FFL motifs over the years, providing significant insights into their functions and mechanisms [19,21,25–28]. Several theoretical analyses of FFLs within the stochastic framework have examined the functional aspects of FFL motifs in noise propagation [17,29–32].

Although FFLs are common in transcription networks, understanding their isolated behavior in terms of noise performance still needs to be completed, particularly in the context of interconnected circuits [10]. Hence, in addition to studying isolated FFLs, it is also important to investigate the behavior of interconnected and overlapping FFLs to gain a more comprehensive understanding of their role in noise performances. In the present manuscript, we consider a multi-output FFL (MOFFL), a higher derivative of FFLs (see Fig. 1a), which is formed by the lateral combination of multiple FFLs having the same input (X) and intermediate (Y) nodes but with multiple outputs (Z_1_, Z_2_, …) [18,22,33]. The MOFFL was found to be the most common generalization of FFL and occurs more frequently in the transcription networks of bacteria and yeast [33]. In *E. coli* transcription network, the maltose utilization system is an example of a MOFFL [33]. Moreover, MOFFL structures are also found in the transcription network of flagellum biosynthesis, where the architecture precisely controls the temporal order in which the motor and filament genes are activated [25,34]. Many other examples of biological model systems of *E. coli* and *S. cerevisiae* are found in which MOFFL structure occurs significantly (see Table. II and III in [33]). We note that the neural network of *Caenorhabditis elegans* predominantly displays a multi-input FFL where more than one input node and an intermediate node regulate the expression of a single output. The MOFFL structure has very few occurrences in this neural network [33].

**Figure 1:**
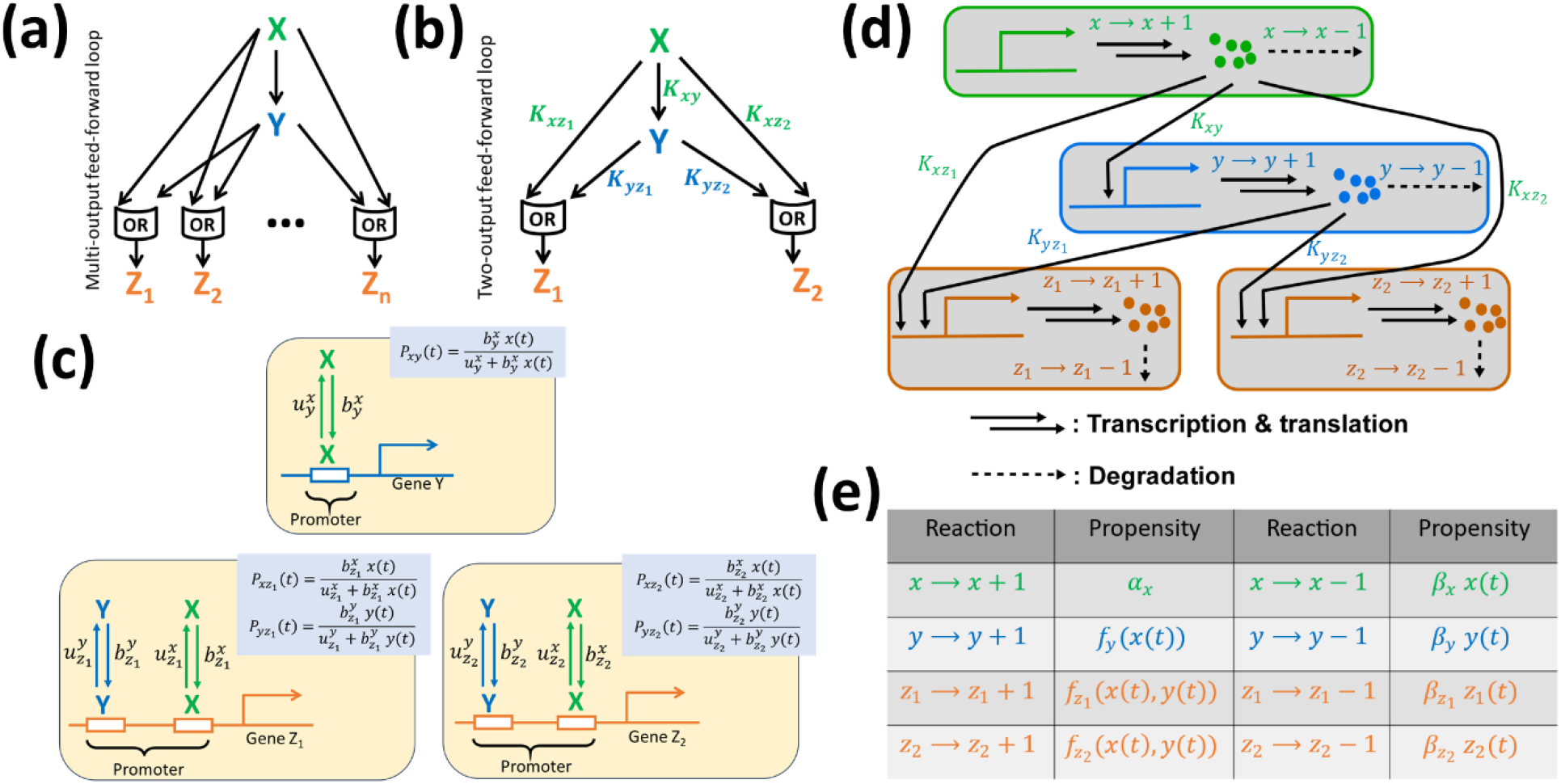
Schematics, molecular interactions and propensities of TOFFL. **(a)** A multi-output feed-forward loop (MOFFL) model with X and Y representing the master and secondary transcription factors (TFs), respectively, and Z_i_ indicating the genes co-regulated by X and Y. **(b)** The two-output feed-forward loop (TOFFL) acts as the fundamental unit of the MOFFL. **(c)** Depiction of TFs binding and unbinding to gene promoters, highlighting the resulting promoter occupancies. **(d)** Detailed kinetic steps of TOFFL and **(e)** corresponding propensities used in numerical simulation using Gillespie’s algorithm [55,56]. The explicit forms of the functions involved in (e) are outlined in Table 1.

In MOFFL, X, the master TF regulates the expression of secondary TF Y, and both jointly co-regulate the set of genes Z_i_ ∈ { Z_1_, Z_2_, Z_3_, ⋯} (Fig. 1a). The co-regulation results in a spectrum of co-expression patterns exhibited by the set of genes Z_i_ depending on the nature and strength of regulations of X and Y. We note that the co-expression of multiple genes refers to the genes being expressed together due to shared TF(s) across diverse conditions. Co-expression with diverse expression patterns highlights that genes can be co-expressed even if they are activated at different times or have varying expression levels. Co-expression data are used to build networks where genes exhibit correlation based on their co-expression patterns [35]. This helps identify groups of genes that likely function together [35,36]. We are interested in the co-expression patterns of genes Z_i_ in MOFFL and its dependence on TFs X and Y regulatory factors. In a recent study, the multi-output incoherent FFL type 1 in the quorum sensing network of *Pseudomonas aeruginosa* was investigated, demonstrating its phenotypic plasticity and robustness in key regulatory factors [37].

**Table 1:**
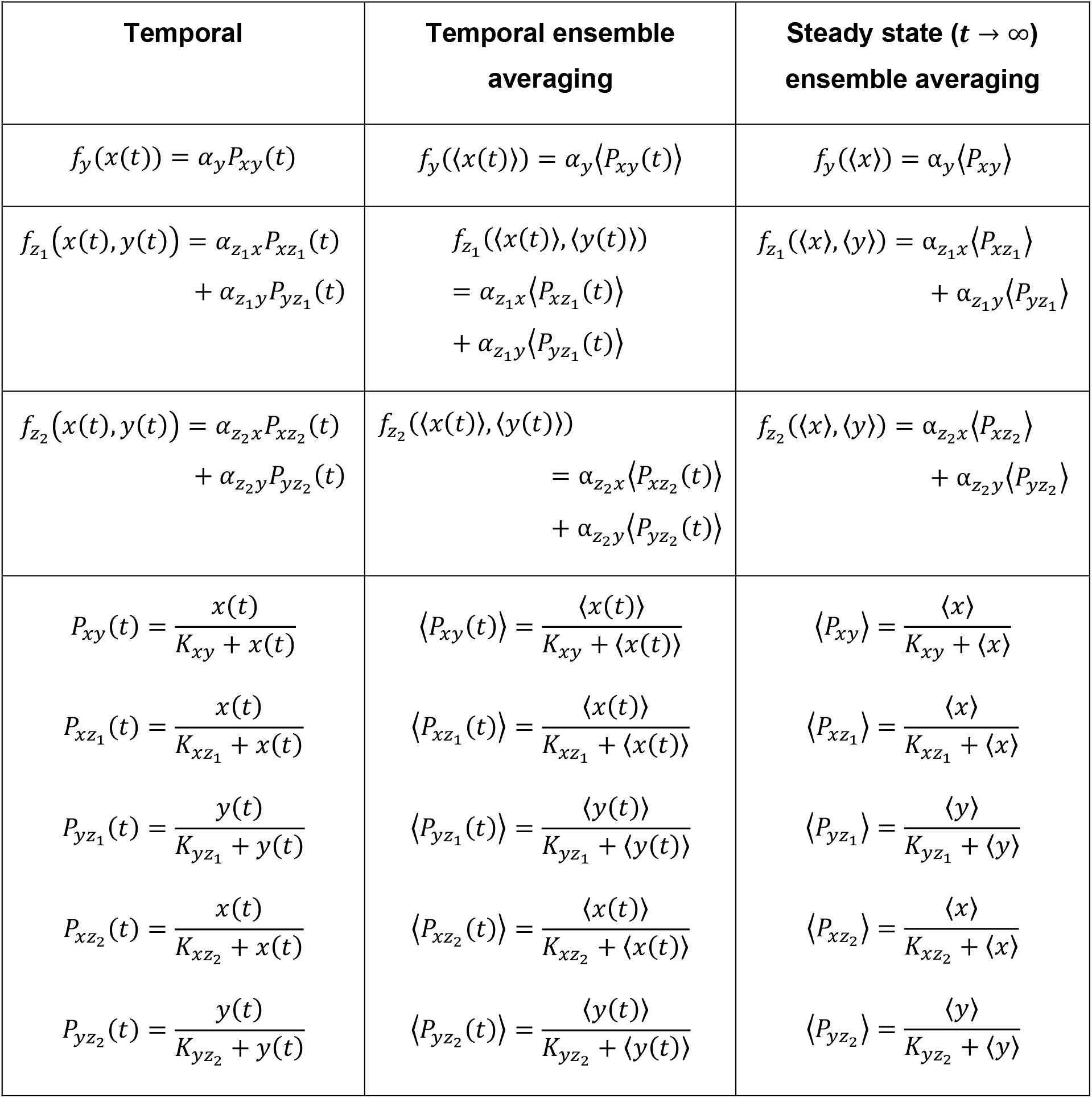
Expressions of synthesis rates at different scenarios. The first column describes how the synthesis rate changes over time in a single realization. The second column shows the average synthesis rate over time by looking at multiple realizations. The third column gives the synthesis rate when the dynamics of the components X, Y, Z_1_, and Z_2_ reach a steady state. The parameter *K*_*i*j_ refers to the activation coefficient, measured in *molecules. V*^−1^. This value essentially sets a threshold level for component *i*. If the concentration of component *i* reaches or surpasses *K*_*i*j_, it significantly boosts the production of component *j* [9]. ⟨⋯ ⟩ represent the averaging over many realizations, providing a population-wide perspective.

We categorize co-expression into symmetric and asymmetric gene expressions. We note that these categories are determined by analyzing protein production profiles, which serve as a proxy for gene expression levels. By symmetric expression, we refer to genes that are activated simultaneously and exhibit equal expression levels across various conditions. In contrast, asymmetric expression pertains to genes that follow similar expression trends but differ in their expression strength and activation timing, indicating a delay in activation. This difference in expression strength can arise due to variations in transcription factor binding affinities, post-transcriptional regulatory mechanisms, or combinatorial control by additional factors. Understanding the biological significance of these co-expression patterns is critical. Symmetric expression may be crucial for processes requiring synchronized activation, whereas, asymmetric expression might allow the cell to be more adaptive to differential environmental signals. By examining these differences, one understands how genes with similar functions are precisely regulated for specific tasks, shedding light on the complex choreography of gene expression that governs cellular responses. In the context of asymmetric gene expression, we refer to a recent study by Ali *et al*. [38], which exemplifies asymmetric co-expression with a negative single-input module, where the transcription factor gene exhibits distinct regulatory asymmetry compared to its target gene.

A critical gap exists in understanding how MOFFLs, with their interconnected regulatory features, influence the inherent noise in gene co-expression. This noise can be beneficial and detrimental [15,24,39,40], so deciphering how MOFFL architecture and regulatory factors influence noise dynamics is crucial. Compared to studying isolated FFLs, investigating MOFFLs provides a more system-level understanding, reflecting the interconnected nature of biological circuits [10].

The present study addresses this gap by investigating the relationship between MOFFL architecture, co-expression patterns (including symmetric and asymmetric), and noise dynamics. We employ a stochastic modeling approach to explore how the binding affinities of the TFs (X and Y) modulate the co-expression and noise properties of MOFFL. Our analysis could guide the development of strategies for manipulating gene expression patterns through targeted modulation of TF binding affinities.

The manuscript is arranged in the following order. In Sec. II and III, we discuss the model and the mathematical aspects of quantifying statistical moments under the stochastic framework. Sec. IV A presents the analysis of the co-expression patterns and their emergence. Sec. IV B addresses the noise dynamics while considering the co-expression of target downstream genes. In Sec. IV C and D, we focus on the coordinated behavior of the output proteins at a particular time and across different time scales. We conclude our study in Sec. V.

## II. MODEL

We model the two-output variant of MOFFL structure, called two-output feed-forward loop (TOFFL). In TOFFL, the master regulator X and the secondary regulator Y regulate the expression of genes Z_1_ and Z_2_ (Fig. 1b). The reduction in model complexity is motivated by recognizing that the TOFFL represents the basic building block observed in the MOFFL structure [22]. This recurring pattern suggests that understanding the dynamics of a TOFFL can provide valuable insights into the more complex regulatory mechanisms of MOFFL. By focusing on this reduced model, we aim to elucidate the fundamental principles of noise propagation underlying the regulatory dynamics of transcription networks in a more tractable manner.

In the TOFFL framework (Fig. 1b), we depict the key players X, Y, and Z_i_ as stochastic variables. This framework enables us to model the system’s dynamic behavior at the molecular level. The input activator, X, is assumed to be synthesized constitutively, i.e., its production occurs independently of other regulatory molecules. However, the master regulator X is typically influenced by various environmental signals that bring fluctuations in X. In contrast, the production of the downstream activator, Y, is controlled by the binding of X to the promoter of gene Y. Again, the expression of gene Z_i_ is regulated by binding X and Y to the respective promoters of gene Z_i_. We note that the binding-unbinding processes of a TF to the promoter of the target downstream gene occur much faster than the time scale of protein fluctuations [9,41]. Keeping this in mind, we simplify the binding-unbinding events as a deterministic quantity [42] called promoter occupancy, which captures the average likelihood of a regulator being bound to the promoter of the downstream gene. The promoter occupancy is then incorporated into the equation for the production rate of downstream proteins. This simplification reduces model complexity while preserving its ability to capture regulatory behavior.

To further enhance analytical tractability, we employ a coarse-grained approach for multi-step molecular processes like transcription and translation [42]. These are grouped into a single, effective step governed by a synthesis function of the respective protein (see Fig. 1d, e). This function incorporates the rate constant and the upstream TF’s promoter occupancy (see Table 1). It captures the net effect of the regulatory interactions on protein production without explicitly considering the steps of transcription and translation.

The temporal evolution of X, Y, Z_1_, and Z_2_ are presented by stochastic differential equations (SDEs) using Langevin approximation [43]. The production and degradation of the master regulator X are modeled as a simple birth-death process following Poisson kinetics to mimic the constitutive growth of X. On the other hand, the X-mediated production of Y is accounted for by the synthesis function *f*_*y*_(*x*(*t*)), while the X and Y-mediated production of Z_i_ is expressed by the function 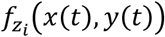. The degradation steps of Y and Z_i_ are modeled using linear functions. The respective SDEs are written as,

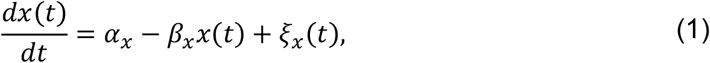

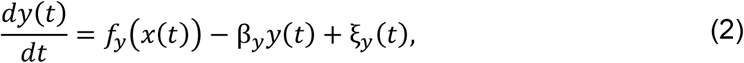

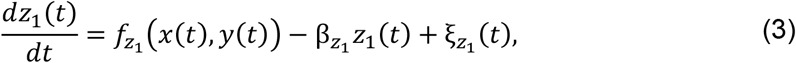

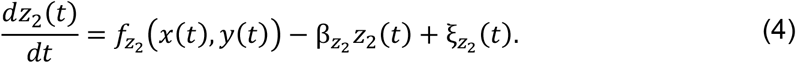

The above set of differential equations is constructed by adding a Langevin random force with the mean dynamics of the components [44–48]. The additive random force (noise) ξ in the above set of equations is assumed to follow Gaussian distribution with the statistics, ⟨ξ_*i*_(*t*)⟩ =0, and ⟨ξ_*i*_ (*t*)ξ_*j*_ (*t*^’^) ⟩ = [*f*_*i*_(⟨⋯ ⟩) + β_*i*_ ⟨*i*(*t*)⟩]δ_*ij*_ δ(*t* − t^’^), where ⟨⋯ ⟩ refer to the ensemble averaging over many realizations and *i* ∈ {*x, y, z*_1_, *z*_2_}. In the definition of noise strength of the variable X, *f*_*i*_(⟨⋯ ⟩) = α_*x*_. The nature of noise correlations suggests that the noise processes are statistically independent with zero mean and zero temporal correlation [12,49–53]. In Eqs. (1-4), *x, y, z*_1_, and *z*_2_ refer to the copy number of the X, Y, Z_1_, and Z_2_, respectively, expressed in *molecules. V*^−1^. In the above equations, 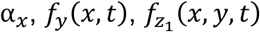 and 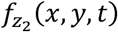 accounts for the synthesis rates of X, Y, Z_1_, and Z_2_, respectively, with unit *molecules. V*^−1^. *min*^−1^. On the other hand, β_*i*_ represents the degradation rate constant of the component *i* with unit *min*^−1^. We note that the TFs X and Y are considered to operate in a manner akin to an OR-gate (also known as a SUM-gate) while regulating Z_i_. The OR gate, here, implies that the presence of either X or Y is adequate to surpass the expression threshold of Z_i_. The explicit forms of the synthesis rates regarding the promoter occupancy are tabulated in Table 1.

The binding and unbinding processes of TFs are illustrated in Fig. 1c. The corresponding rate constants for binding and unbinding events are 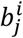 and 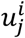, respectively. Here, *i* represents the specific TF, while *j* corresponds to the gene whose promoter is available for binding (*i* ∈ {*x, y*} and *j* ∈ {*y, z*_1_, *z*_2_}). Given that the time scale of binding-unbinding processes is significantly faster than the time scales β_*i*_, these processes rapidly reach equilibrium. Consequently, the promoter occupancy is determined by the expression 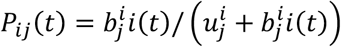 (Fig. 1c). By defining the binding coefficient as 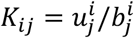 [54], the promoter occupancy can be rewritten as *P*_*ij*_ (*t*) = *i*(*t*)/ *K*_*ij*_ + *i*(*t*), which now depends explicitly on binding coefficient, *K*_*ij*_. We note that the activation coefficient *K*_*ij*_ in the expression of promoter occupancy signifies the threshold concentration of *i* required to express *j* maximally. It is described in the unit of copy number, i.e., *molecules. V*^−1^. The corresponding expressions of the promoter occupancy are provided in Table 1.

The promoter occupancy, denoted by *P*_*ij*_ (*t*), reflects the level of TF (*i*(*t*)) binding to the promoter of gene *j* at a specific time *t*. Since the copies of the upstream TF, *i*(*t*), vary over time, *P*_*ij*_ (*t*) will also vary accordingly. If one considers the average concentration of the TF, denoted by ⟨*i*(*t*)⟩, at time *t*, one can use the expression of average promoter occupancy, ⟨ *P*_*ij*_ (*t*) ⟩. When the protein production-degradation events reach a steady state where protein production is constant, the TF concentration can also be represented by a single, steady state value, ⟨*i*⟩. In this scenario of steady state system dynamics, the promoter occupancy becomes a constant value, ⟨*P*_*ij*_⟩ (see Fig. 2c).

**Figure 2:**
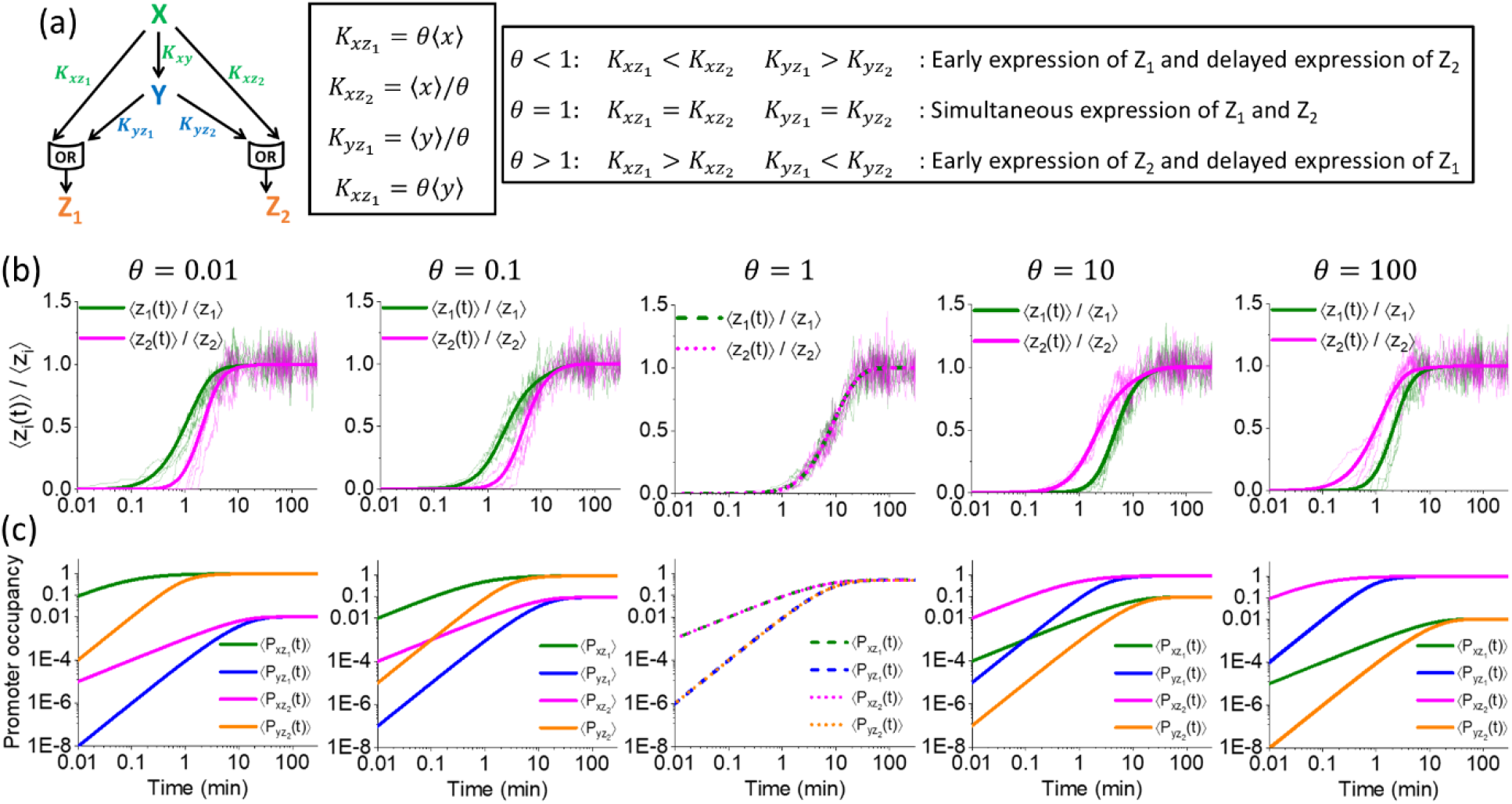
DBA regulates symmetric and asymmetric expression patterns of the output proteins. **(a)** Introduction of the differential binding affinity, θ, and its relation with the activation coefficients (*K*_*ij*_). **(b)** Temporal evolution of normalized average copy number of Z_1_ and Z_2_ for various values of θ. The normalization is achieved by dividing the mean copy number of Z_i_ at time *t* (⟨*z*_*i*_(*t*)⟩) with the corresponding steady state copy number, represented by ⟨*z*_*i*_⟩. The mean copy number ⟨*z*_*i*_(*t*)⟩ are evaluated by solving Eqs. (9-12) numerically. The corresponding steady state values are set at ⟨*z*_1_⟩ = ⟨*z*_2_⟩ = 100 *molecules*/*V*. The normalized mean profiles are drawn as solid lines (theoretical) on top of some trajectories (faint and fluctuating lines) generated numerically using Gillespie’s algorithm [55,56]. Five sample trajectories are generated for Z_1_ and Z_2_ each. The other parameters used to generate the solid lines and trajectories are ⟨*x*⟩ = ⟨*y*⟩ = 100 *molecules* /*V*, β_*x*_ = β_*y*_ = 0.1 *min*^−1^, and 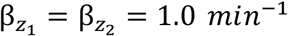. The synthesis rate constants α_*i*_ are set using Eqs. (5-8). **(c)** The temporal profile of average promoter occupancy, as listed in Table 1.

In summary, promoter occupancy is not a fixed value. It depends on the current state of the system. The occupancy will vary during the initial, transient phase of gene expression. However, once the system reaches a steady state, the promoter occupancy becomes a constant value reflecting the average TF level bound to the promoter.

## III. METHOD

Using Eqs. (1-4), we have the steady state copy number of X, Y, Z_1_, and Z_2_,

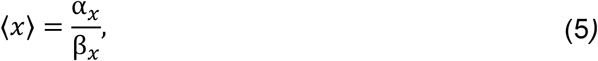

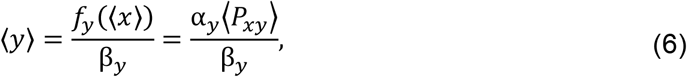

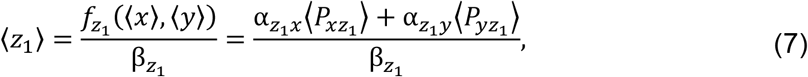

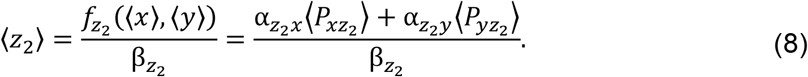

We utilize Eqs. (5-8) to evaluate the synthesis rate constants, α’s. In this regard, we assume that the maximal production rates of Z_1_ from both X and Y are equal, and similarly for Z_2_. Mathematically we express these assumptions as 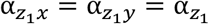 and 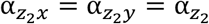. The expressions of the synthesis rate constants are 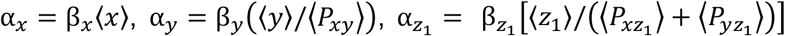, and 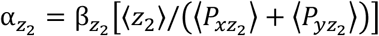. Throughout the manuscript, we employ these expressions of α’s to compute various measures for the TOFFL.

Since Eqs. (1-4) are nonlinear, obtaining analytical solutions at the transient limit is challenging. Therefore, we resort to numerical simulation using the Gillespie algorithm [55,56] to compute the first and second moments in the transient regime. To perform the simulation, we present the detailed kinetic steps and corresponding propensities involved in TOFFL in Fig. 1d, e. However, a steady state solution of Eqs. (1-4) can be derived to calculate the second moments. To obtain analytical expressions for the second moments at steady state, we use the Lyapunov equation for the covariance matrix [12,52,57]. We refer to the *Appendix A* for a detailed step-wise derivation of the steady state second moments.

## IV. RESULTS

### A. Differential binding affinity and asymmetric gene expression

In TOFFL, activation coefficients (*K*_*ij*_) play critical roles in orchestrating the differential activation of genes Z_1_ and Z_2_. These coefficients essentially act as thresholds for gene expression, where a lower *K*_*ij*_ value signifies a lower concentration of the regulatory molecule (TF) needed for robust gene activation [9]. The master regulator, X, in the TOFFL, influences three genes, Y, Z_1_, and Z_2_, through activation coefficients 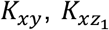, and 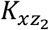, respectively. Similarly, the secondary regulator, Y regulates Z_1_ and Z_2_ with coefficients 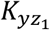 and 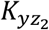.

We, now, introduce a coefficient θ representing differential binding affinity (DBA), which alters the activation coefficients *K*_*ij*_ to establish a hierarchy, leading to the asymmetric expression of genes Z_1_ and Z_2_. Specifically, we define the following relations: 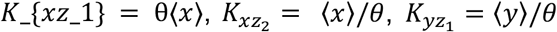, and 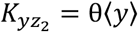. It is important to note that 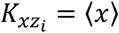 and 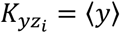 correspond to the half-maximal expression of Z_i_ by X and Y, respectively. Therefore, the introduction of θ effectively alters the half-maximal regulation, thereby influencing the relative binding strength of TFs to the promoters of the genes Z_1_ and Z_2_.

When 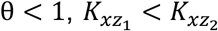 and 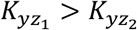, indicating that Z_1_ is activated before Z_2_ (Fig. 2b). This delay correlates with promoter occupancy (Fig. 2c), where 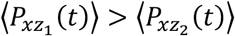 suggesting stronger binding of X to the promoter of gene Z_1_, leading to preferential activation. This reinforces the notion that lower *KK*_*ij*_ values result in faster activation due to stronger promoter occupancy. On the other hand, when 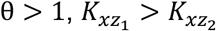 and 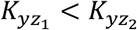, indicating the activation of Z_2_ precedes Z_1_ (Fig. 2b) and the promoter occupancy follows the opposite trend as observed in the case of θ< 1 (Fig. 2c). Notably, as the deviation of θ from 1 increase (θ< 1 or θ> 1), the separation between the temporal activation profiles of Z_1_ and Z_2_ becomes more pronounced (Fig. 2b). In other words, a larger deviation from θ= 1 leads to a greater asymmetry observed in the expression pattern of these genes. However, when 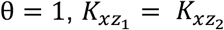 and 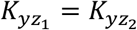, indicating simultaneous expression of the two genes Z_1_ and Z_2_, resulting in identical and symmetric temporal dynamics (Fig. 2b, c). In essence, θ captures the differential affinity of the TFs to the promoters of gene Z_1_ and Z_2_. A value of θ= 1 indicates equal binding affinity, while deviations from 1 reflect asymmetry in TF’s binding, resulting in asymmetric gene expression. We note that at *t* ≈ 100 min, the dynamics of both Z_1_ and Z_2_ reach the steady state. In the rest of the manuscript, the steady state profiles of noise and correlation are measured after *t* = 100 min, where the dynamics of Z_1_ and Z_2_ reach the steady state.

The bold lines in Fig. 2b represent the normalized average behavior of Z_i_ obtained from theoretical calculations. However, the inherent fluctuations are captured in the faint, fluctuating lines in Fig. 2b. These lines represent individual trajectories obtained in stochastic simulations using Gillespie’s algorithm [55,56].

The temporal dynamics presented as bold lines in Fig. 2b are derived by computing the ensemble averages of Eqs. (1-4) at time *t*, resulting in the following coupled differential equations,

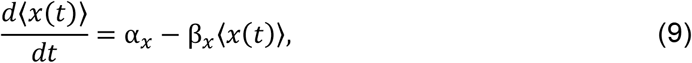

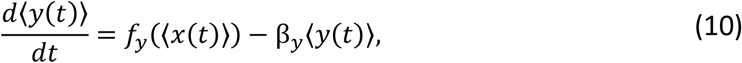

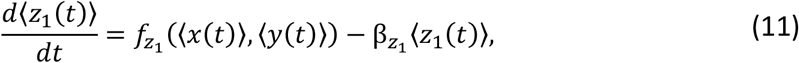

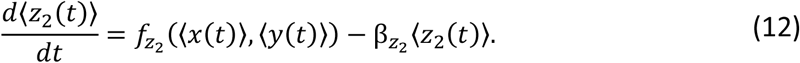

The coupled Eqs. (9-12) are solved numerically as the analytical solution is complex due to its non-linear nature. In Fig. 2, we use the following relaxation rate parameters 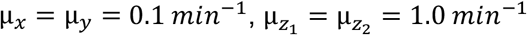. The steady state copy number of each component is taken equal, i.e., ⟨*x*⟩ = ⟨*y*⟩ = ⟨*z*_1_⟩ = ⟨*z*_2_⟩ = 100 *molecules*.*V*^−1^, for better comparing the results on equal footing. In addition, we use the half-maximal expression of Y by X where *K*_*xy*_ = ⟨*x*⟩. Due to the presence of asymmetry in gene expression, we expect the noise properties of the system’s components to be altered at both transient and steady state levels. This motivates us to explore how the DBA, θ modulates noise.

### B. Emergence of hierarchy in noise

One common way to quantify the noise is to compute the Fano factor (FF). The FF of a random variable is defined as the ratio of variance to the mean. In the context of TOFFL, we compute the temporal FF of Z_1_ and Z_2_ using the definition, 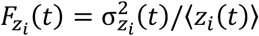 where 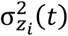 and ⟨*z*_*i*_(*t*)⟩ refer to the temporal variance and mean of the output proteins. Since the temporal solution of the non-linear equations (1-4) are analytically intractable, we employ the numerical method of Gillespie’s algorithm [55,56]. The purpose of FF is to understand the dynamics of the system in comparison to the Poisson scale (*F* = 1). When examining the effect of the DBA, θon the FF, distinct behaviours emerge. For θ< 1, Z_1_ is activated before Z_2_ (Fig. 2b) and Z_2_ exhibits a higher FF compared to Z_1_ (Fig. 3a). Conversely, when θ> 1, Z_2_ is activated prior to Z_1_ (Fig. 2b) and the FF of Z_1_ surpasses that of Z_2_ (Fig. 3a). These observations suggest that genes with delayed expression tend to have higher FF, while those activated earlier demonstrate a lower FF. When θ= 1, both the proteins show a collating nature in the FF profile (Fig. 3a). In addition, as the deviation of θ from 1 increase, the separation between the FF profiles of Z_1_ and Z_2_ increases as had been observed in the expression profiles (Fig. 2b).

**Figure 3:**
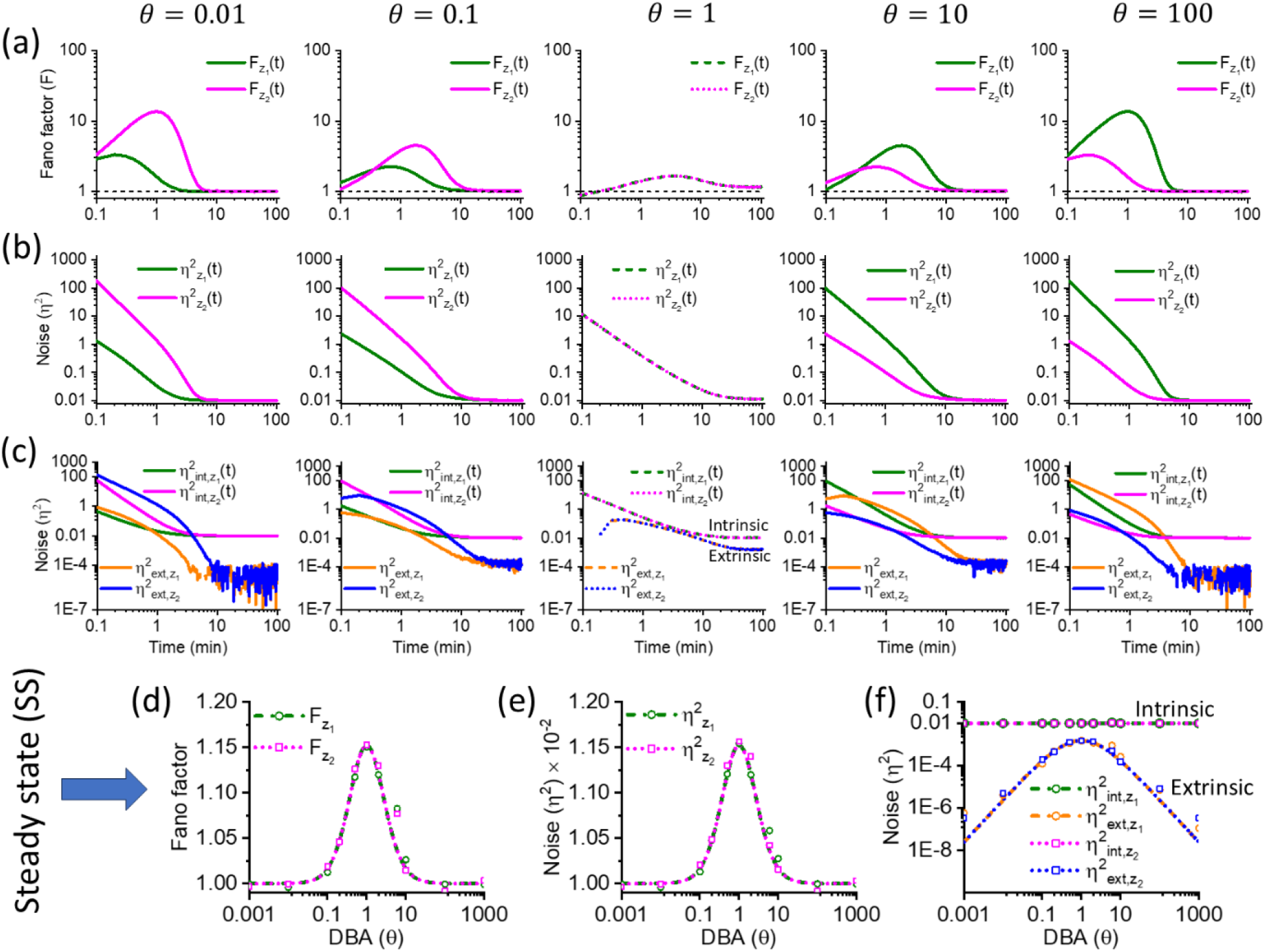
Asymmetric expression induces temporal hierarchy in noise. **(a)** Temporal profiles of Fano factor, **(b)** noise measured by square of CV, and **(c)** intrinsic and extrinsic noise components for various values of θ. The profiles (a), (b), and (c) are generated by simulating the dynamics using Gillespie’s algorithm [55,56]. Profiles of **(d)** Fano factor, **(e)** noise, and **(f)** intrinsic and extrinsic noise components as a function of θ at steady state. The lines in (d), (e), and (f) are generated using analytical expressions derived at steady state (see *Appendix A*). The points in the same profiles are due to stochastic simulation. The parameters used to generate the figures (a)-(f) are, ⟨*x*⟩ = ⟨*y*⟩ = 100 *molecules* /*V*, ⟨*z*_1_⟩ = ⟨*z*_2_⟩ = 100 *molecules* /*V*, β_*x*_ = β_*y*_ = 0.1 *min*^−1^, and 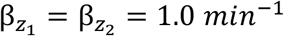. The synthesis rate constants α_*ii*_ are set using Eqs. (5-8).

Another characteristic of the FF profiles is that during the transient regime, FF exceeds 1 for Z_1_ and Z_2_ at any θ value. However, for θ< 1 and θ> 1, FF converges to unity as the outputs approach a steady state (FF ≈ 1). This indicates that during the transient regime, a higher level of variability is associated with both proteins compared to the Poisson process. Notably, FF = 1 signifies that the noise is solely attributed to the intrinsic birth-death processes involved with the proteins, indicating purely intrinsic noise. Conversely, FF > 1 suggests the presence of significant external noise sources, such as the noise from X and Y, which amplify the noise level of each output protein beyond their intrinsic level at the transient limit. These external noises are referred to as extrinsic noises. Once Z_1_ and Z_2_ reach a steady state, the impact of external noise sources becomes insignificant, resulting in FF ≈ 1, representing the typical Poisson-like signature. One point should be noted that the scenario for θ= 1 is different where FF still has a slightly larger value than 1 at steady state. This indicates that the extrinsic noise sources still inject some degree of noise into the steady state noise level of Z_1_ and Z_2_ at θ= 1.

We, further, analyzed the square of the coefficient of variation (*CV*^2^), of Z_1_ and Z_2_, defined as 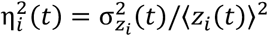. We opt for *CV*^2^ as the measure is dimensionless and provides the relative variability in the expression levels compared to the mean, independent of the absolute concentration. As expected, the *CV*^2^ analysis aligns perfectly with the FF observations, with the earlier activated gene exhibiting lower noise compared to the delayed one (Fig. 3b). Moreover, we decompose the total noise into intrinsic 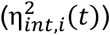 and extrinsic 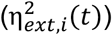 parts, which hold the relation [14,16],

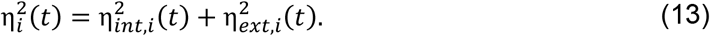

We note here that the intrinsic noise is solely attributed to the inherent birth-death dynamics of the respective proteins, while extrinsic noise accounts for the fluctuation injected into the protein level due to upstream regulators X and Y. To obtain the expression of intrinsic noise, 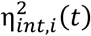 we make use of the fact that when there are no extrinsic factors (X and Y), the intrinsic noise reflects the total noise which arises due to the simple birth-death dynamics of Z_i_. As a result, the dynamics follow Poisson distribution and FF = 1, which signifies that the variance of Z_i_ equals mean, 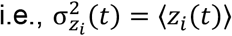. In this case, the total noise and, hence, the intrinsic noise would become,

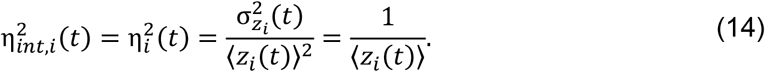

When X and Y are present, the extrinsic noise can be evaluated as 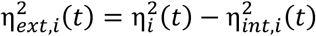 (Eq. (13)). For θ< 1 and θ> 1, the extrinsic noise of Z_1_ and Z_2_ at the transient regime is comparable to the corresponding intrinsic noise. At the same time, it becomes insignificant as the protein dynamics reach the steady state where only intrinsic noises are dominant (Fig. 3c). At θ= 1, the extrinsic noise remains significant at transient and steady state regimes and cannot be disregarded (Fig. 3c).

To connect the noise decomposition with FF, we use the definition of FF, *CV*^2^, and Eqs. (13-14) to write (see *Appendix B*),

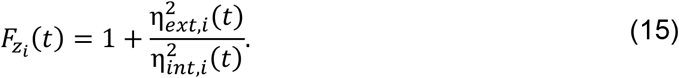

Eq. (15) indicates that when there are no external sources of noise 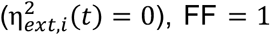. However, the extrinsic noise causes the FF to exceed 1 in the transient phase. This perfectly aligns with the observation in temporal FF profiles. At steady state, 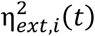 is negligible (Fig. 3c) and results in FF → 1 (Fig. 3a).

To explore the steady state noise behaviors of the system, we analyze the system separately in the steady state regime. At steady state, the FF of both Z_1_ and Z_2_ converges. This characteristic is also evident in Fig. 3a. When θ= 1, maximum FF is achieved, while deviation from it results in a decrement of FF, finally converging to FF = 1 (Fig. 3d). The steady state FF is defined as 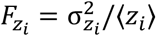 where 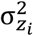 and ⟨*z*_*i*_ ⟩ refer to the steady state variance and mean of Z_i_, respectively. Similarly, the steady state noise is quantified by 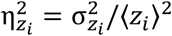. We note that Eqs. (14-15) are also applicable to the steady state. To obtain the expressions of steady state FF and noise, we refer to *Appendix A*. The noise profiles of Z_1_ and Z_2_ show a similar nature as observed for FF (Fig. 3e). Further decomposing the steady state noise into intrinsic and extrinsic noise reveals that when θ< 1 or θ> 1, the magnitude of extrinsic noise is insignificant, implying the dominant role of intrinsic noise (Fig. 3f). However, at θ= 1, the extrinsic noise is significant enough to contribute to the total noise. We refer to *Appendix A* for steady state noise decomposition.

Our analysis reveals a relationship between the timing of gene activation and the noise levels in the system. We find that asymmetric temporal activation order is inversely related to the order of noise levels in the transient limit. Genes activated early in the process experience lower noise levels, attributed to reduced intrinsic and extrinsic fluctuations associated with gene expression. Conversely, genes expressed with a delay exhibit higher noise levels, characterized by elevated intrinsic and extrinsic noise levels. This finding highlights the intricate balance between temporal regulation and noise modulation in gene expression.

### C. Instantaneous correlation between output protein’s expression patterns

We now examine the collective behavior of Z_1_ and Z_2_, focusing on how they interact during transient and steady state phases of gene expression. To understand their interplay, we employ two key metrics, Pearson’s correlation coefficient and mutual information (MI) between Z_1_ and Z_2_ at a single snapshot in time, *t*. Pearson’s correlation coefficient is defined as 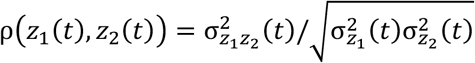, where 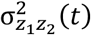 refers to the temporal covariance computed numerically (see Fig. 1d, e which outlined the kinetics used in simulation). Since the system variables are Gaussian distributed, the MI between Z_1_ and Z_2_ is defined as, i.e., *I*(*z*_1_(*t*); *z*_2_(*t*)) = −(1/2) log_2_ (1 − ρ^2^ (*z*_1_(*t*), *z*_2_(*t*))) [58]. The correlation coefficient, ρ(*z*_1_(*t*), *z*_2_(*t*)) quantifies the direction and strength of the relationship between Z_1_ and Z_2_ at a specific time, *t*. A positive correlation (ρ(*z*_1_(*t*), *z*_2_(*t*)) > 0) indicates that their expression levels tend to move together, suggesting coherence or coordinated activity. Conversely, a negative correlation (ρ(*z*_1_(*t*), *z*_2_(*t*)) < 0) reveals an incoherent relationship, where one gene’s expression increases as the other decreases. On the other hand, the MI, *I*(*z*_1_(*t*); *z*_2_(*t*)) captures the information sharing between Z_1_ and Z_2_ at a specific time point, *t*. The idea behind measuring MI is to capture how much information they share and how this information correlates with the observed temporal asymmetry in gene expression. A higher MI value indicates a richer exchange of information. For example, if the expression level of Z_1_ provides a good prediction of the expression level of Z_2_ (and vice versa) at a particular time *t*, the MI will be high. Conversely, a low MI suggests limited information exchange between the two genes at that specific time. Unlike the correlation coefficient, MI doesn’t tell us the direction of the relationship (coordinated or opposite behaviour) but focuses on the amount of information they share.

We observe non-monotonic patterns in both the correlation and mutual information (MI) over time (Fig. 4a, b). These metrics peak during the transient phase and decrease as the system approaches the steady state. The timing and magnitude of these peaks vary depending on θ. For significant deviations from 1 (e.g., θ= 0.01 and 100), the correlation and MI peak earlier, albeit with reduced magnitudes. Conversely, at θ= 1, the peak occurs later, and its magnitude is higher. Simultaneous expression of Z_1_ and Z_2_ (θ= 1) leads to maximal information sharing and positive correlation, indicating highly coordinated dynamics. However, enforcing temporal asymmetry in expression (θ≠ 1) requires either θ> 1 or < 1. In such cases, Z_1_ and Z_2_ exhibit less coordinated behavior, with lower information sharing and positive correlation than θ= 1. The steady state behaviour of correlation and MI as a function of θ is depicted in Fig. 4c, d, showing a higher degree of correlation between the genes at θ= 1, which diminishes upon deviation from 1. We note that the definitions of Pearson’s correlation coefficient and MI also apply to the steady state measure, given that the corresponding variances and covariance are measured at a steady state (see *Appendix A*).

**Figure 4:**
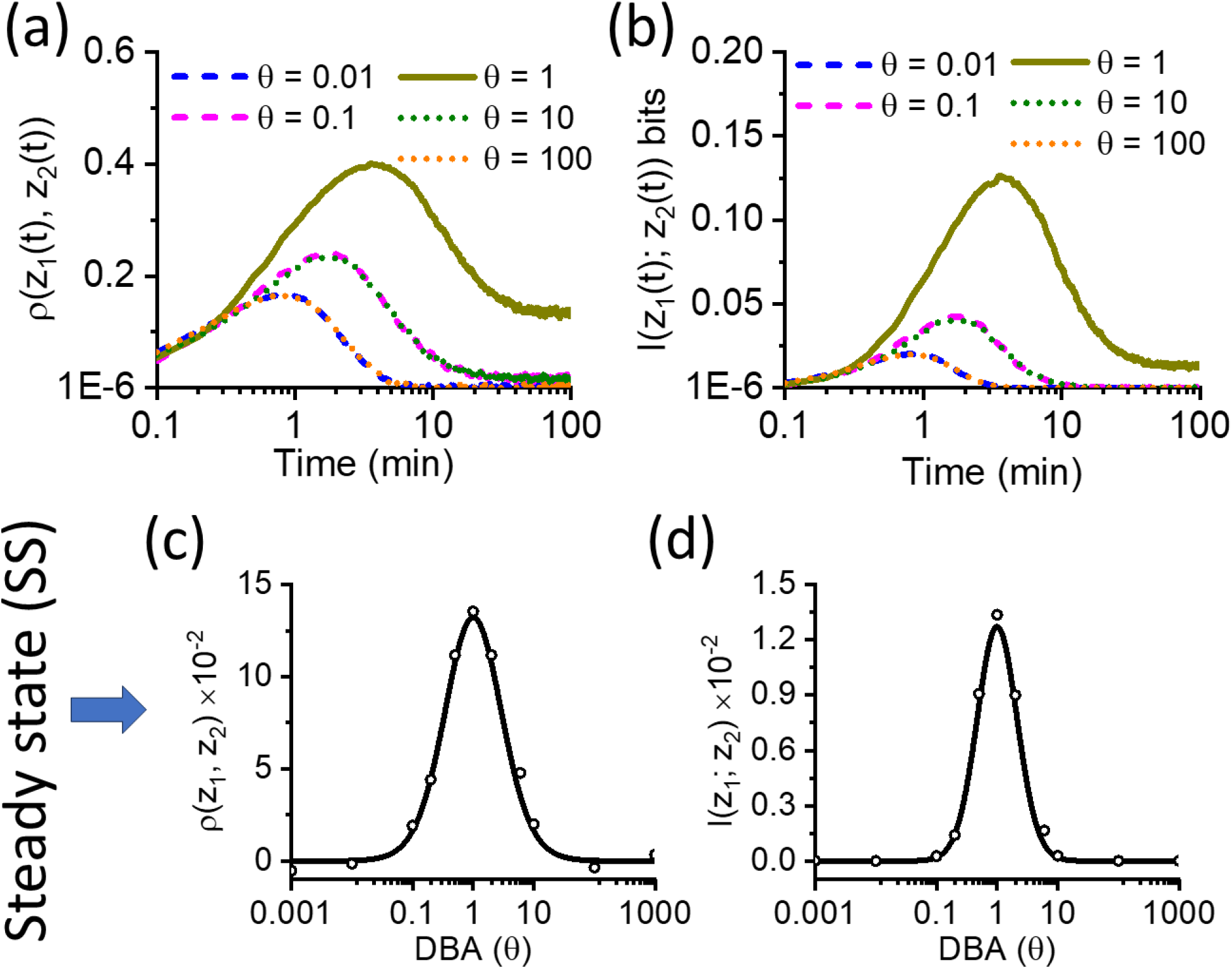
Symmetric and asymmetric expressions modulate the instantaneous correlation and information sharing between the output proteins. Temporal profiles of **(a)** instantaneous correlation coefficient, ρ(*z*_1_(*t*), *z*_2_(*t*)) and **(b)** mutual information, *I*(*z*_1_(*t*); *z*_2_(*t*)) as a function time for various values of θ. Steady state profiles of **(c)** correlation coefficient, ρ(*z*_1_, *z*_2_) and **(d)** mutual information, *I*(*z*_1_; *z*_2_) as a function of θ. The lines are due to stochastic simulation in (a) and (b). In (c) and (d), lines are due to analytical expressions (see *Appendix A*), and the points are generated by stochastic simulation using Gillespie’s algorithm [55,56]. The parameters used to generate the figures (a)-(d) are the same as mentioned in Fig. 3.

The temporal and steady state behavior of correlation and MI can also be explained by considering the noise dynamics of Z_1_ and Z_2_. At the transient phase, both intrinsic and extrinsic noise are prevalent (Fig. 3c) at every value of θ. This extrinsic noise originating from shared regulators X and Y introduces a correlation between the expression levels of Z_1_ and Z_2_ in the transient phase, reflected in the correlation and MI profiles (Fig. 4a, b). However, for θ≠ 1, the extrinsic noise becomes less prominent as the system reaches the steady state (Fig. 3c), and the intrinsic noise, originating from the gene expression process itself, plays the dominant role in this regime. This leads to a decoupling in their expression patterns to a certain extent. With less influence from shared upstream fluctuations and more reliance on specific regulatory cues, Z_1_ and Z_2_ become less reliant on each other’s immediate expression levels for guidance, resulting in a decrease in correlation and information sharing towards the steady state. Interestingly, even though the expression levels of Z_1_ and Z_2_ might appear “merged” at the steady state (Fig. 2a), the communication between them becomes less frequent (Fig. 4c, d).

Analysis of Z_1_ and Z_2_ dynamics reveals that temporal asymmetry in gene expression, governed by DBA θ, influences gene coordination and communication. Simultaneous expression (θ= 1) leads to maximal coordination, while asymmetric expression (θ≠ 1) reduces coordination and inter-protein communication. Loss of communication is detrimental, particularly with prolonged delays (θ≫ 1 or ≪ 1), underscoring the importance of precise temporal regulation of gene expression and cellular function.

### D. Time-delayed correlation between output proteins

This section analyzes the time-delayed correlation and MI between the expression levels of Z_1_ and Z_2_. These measures capture how their expression levels are related across different time points. We first quantify the auto-correlation function (Fig. 5a) for both genes. This function, defined as 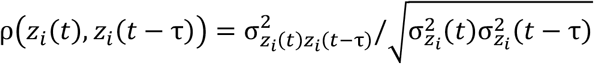, measures how similar the expression level of gene Z_i_ at time *t* is to its expression level at a previous time point *t* − τ. The calculation involves 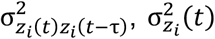, and 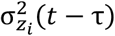, which refer to the covariance between *z*_*i*_ (*t*) and *z*_*i*_(*t* − τ), variance of *z*_*i*_(*t*), and *z*_*i*_ (*t* − τ), respectively. The variances and covariances are computed using simulation and the detailed kinetic steps used in the simulation are outlined in Fig. 1d, e. Similarly, the time-delayed self-MI, defined as *I*(*z*_*i*_(*t*); *z*_*i*_(*t* − τ) = −(1/2) log_2_ (1 − ρ^2^ (*z*_*i*_(*t*), *z*_*i*_(*t* − τ), captures the information shared between the expression level of Z_i_ at time *t* and its own level at time *t* − τ.

**Figure 5:**
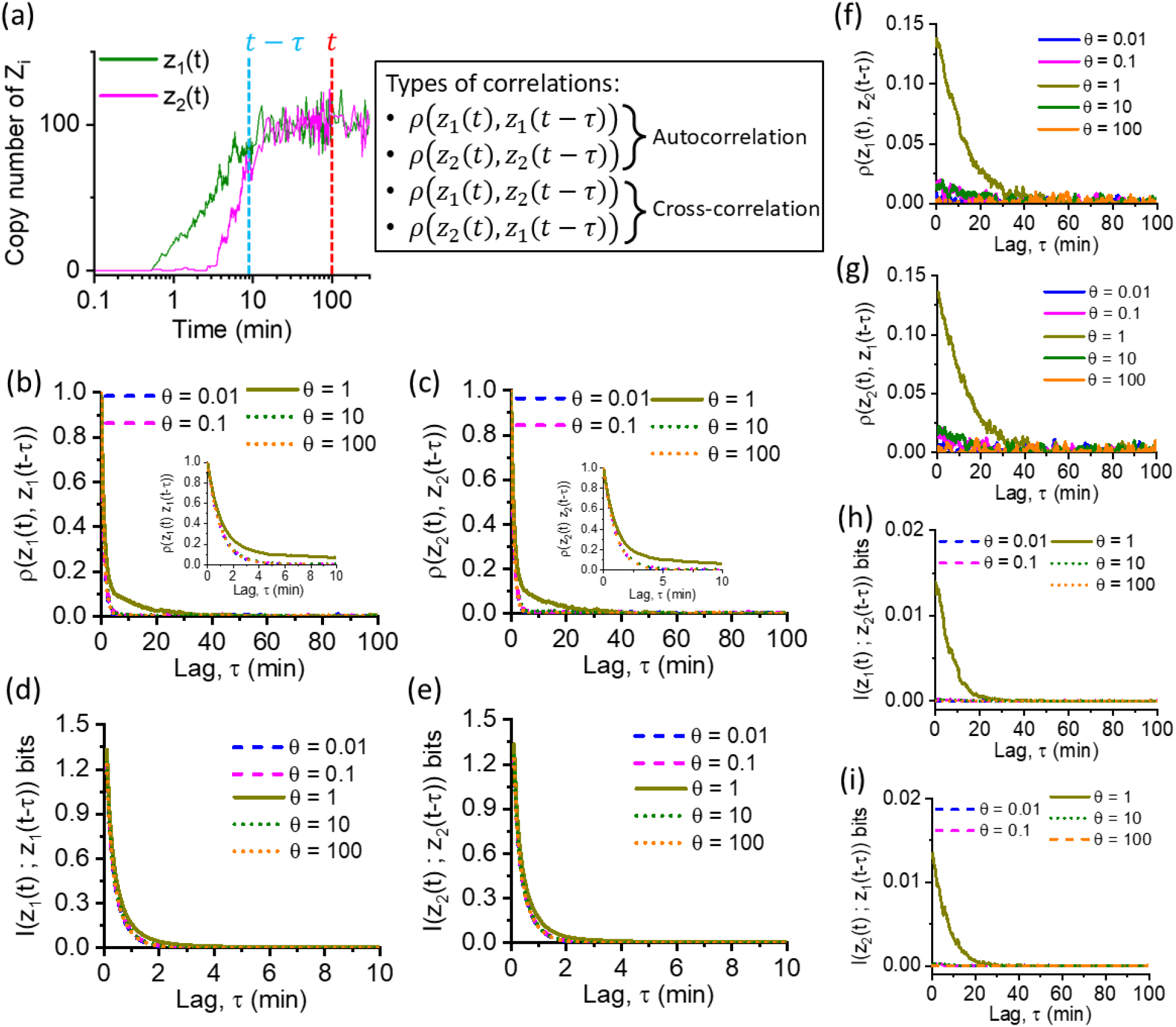
Effect of symmetric/asymmetric expression patterns on time-delayed correlation and information sharing between the output proteins. **(a)** Schematics of various correlations, measured between the output proteins Z_1_ and Z_2_. **(b-c)** Temporal profiles of the auto-correlations, ρ(*z*_1_(*t*), *z*_1_(*t* − τ)) and ρ(*z*_2_(*t*), *z*_2_(*t* − τ)) for various values of θ. **(d-e)** Temporal profiles of the self-mutual information, *I*(*z*_1_(*t*), *z*_1_(*t* − τ)) and *I*(*z*_2_(*t*), *z*_2_(*t* − τ)) for various values of θ. **(f-g)** Temporal profiles of the cross-correlations, ρ(*z*_1_(*t*), *z*_2_(*t* − τ)) and ρ(*z*_2_(*t*), *z*_1_(*t* − τ)) for various values of θ. **(h-i)** Temporal profiles of the cross-mutual information, *I*(*z*_1_(*t*), *z*_2_(*t* − τ)) and *I*(*z*_2_(*t*), *z*_1_(*t* − τ)) for various values of θ. The lines in (b-i) are generated by numerical simulation using Gillespie’s algorithm [55,56]. The parameters used to generate the figures (b)-(i) are the same as mentioned in Fig. 3. While measuring cross-correlations and cross-mutual information, we use *t* = 100 *min*.

The auto-correlation for both Z_1_ (Fig. 5b) and Z_2_ (Fig. 5c) exhibit a similar profile. They show a positive correlation at zero lag (τ = 0) for all θ values, indicating that the expression level of a gene at a specific time tends to be similar to its level at the same time point. The correlation value quickly decays towards zero as the lag (τ) increases. The decay is much faster for θ≠ 1 compared to θ= 1. This decay signifies a weakening relationship between the expression level of Z_i_ at a specific time (*t*) and its level at progressively earlier time points (*t* − τ). The auto-correlation function converging to zero after a few lags suggests that the influence of past expression levels on the current expression becomes negligible after a certain time window. Similarly, the time-delayed self-MI also shows a peak at zero lag and then decreases with increasing lag (Fig. 5d, e). This indicates that the information a gene’s current expression level carries about its past expression levels diminishes as we trace back to the temporal profile. A gene’s current expression level is mainly influenced by its recent past expression and weakens as we consider progressively older expression data.

Next, we investigate the correlation and information sharing between Z_1_ and Z_2_ across different time scales. We quantify this by calculating the cross-correlation coefficient ρ(*z*_*i*_ (*t*), *z*_*j*_(*t* − τ)) (*i, j* ∈ {1,2}) between their expression levels (i.e., *z*_*i*_(*t*) for gene product *i* at time *t* and *z*_*j*_ (*t* − τ) for gene product *j* at a lagged time point *t* − τ). The cross-correlation, defined as 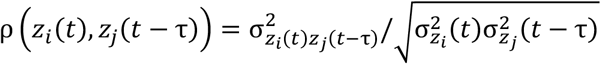 incorporates the covariance 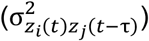 between the expression levels at different times and normalizes it by their variances 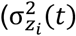 and 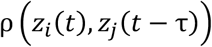 The variances and covariances are computed using stochastic simulation. The metric, ρ (*z*_*i*_(*t*), *z*_*j*_(*t* − τ)), reflects how the expression level of gene *i* at time *t* is related to the expression level of gene *j* at a previous time point *t* − τ. We also calculate the time-delayed cross-MI, defined as *I* (*z*_*i*_ (*t*); *z*_*j*_(*t* − τ) = −(1/2) log_2_ 1 − ρ^2^ (*z*_*i*_(*t*), *z*_*j*_(*t* − τ)), to quantify the information exchange between Z_1_ and Z_2_ across different time lags.

A significant cross-correlation between Z_1_ and Z_2_ is observed only up to a certain lag (τ) and exclusively for the case where θ= 1 (i.e., simultaneous expression) (Fig. 5f, g). This suggests that when the genes are activated simultaneously and symmetrically, their expression levels are correlated across short time windows. However, introducing a temporal asymmetry in gene expression (θ≠ 1) leads to an insignificant cross-correlation (ρ (*z*_*i*_(*t*), *z*_*j*_ (*t* − τ) ≈ 0) across all time scales (Fig. 5f, g). This indicates that when expression is asymmetric, the expression of one gene no longer carries information about the past expression of the other across different time lags.

A similar trend is observed for the time-delayed cross-MI (Fig. 5h, i). When θ= 1, a certain level of information is transferred from the previous expression level of Z_1_ (Z_2_) to the current expression level of Z_2_ (Z_1_). However, this information exchange becomes negligible after a specific lag (τ). This suggests that when the genes are activated simultaneously, there is a limited window for information exchange about their past expression levels.

The analysis of cross-correlation and time-delayed cross-MI highlights the importance of simultaneous expression (θ= 1) for coordinated information exchange between Z_1_ and Z_2_. Introducing an asymmetry disrupts this information flow, leading to a loss of correlation and information sharing across different time scales. Our findings suggest that inter-protein communication fosters a more coordinated (coherent) gene expression pattern. Conversely, asymmetric gene expression leads to incoherence and hinders communication between the genes.

## V. DISCUSSION

This manuscript investigates a recurring structure in transcription networks, the MOFFL [18,22,33]. We simplify the MOFFL structure to its core building block, the TOFFL. To analyze the stochastic behavior of this network, we model the TOFFL dynamics using SDEs within the Langevin framework [43]. A key coefficient θ is introduced to capture the differential binding affinity (DBA) of upstream TFs (X and Y) to the promoters of downstream genes Z_1_ and Z_2_. We quantify the noise of the output protein levels of the TOFFL, both transiently and at steady state, using the Fano Factor (FF) and squared coefficient of variation (*CCVV*^2^). Furthermore, we explore how DBA (θ) influences the coordinated behaviour of the two output proteins, Z_1_ and Z_2_. The coordination is measured through correlation coefficients and mutual information (MI) at specific time points and across various time scales. The key outcomes of our analysis are the following.

The coefficient, θ, significantly influences the expression patterns of downstream genes Z_1_ and Z_2_. When θ= 1, both genes are activated simultaneously with equal strength, resulting in symmetric protein expression. However, when θ deviates from one, asymmetric expression emerges, where one gene is activated earlier than the other. This asymmetry in activation timing directly impacts protein noise levels, with the earlier-activated gene exhibiting lower noise, thus emerging a temporal hierarchy in noise levels between Z_1_ and Z_2_. Notably, when both genes are activated simultaneously (θ= 1), their noise levels are equal. Noise analysis reveals contributions from both intrinsic and extrinsic sources, particularly during the transient phase. While intrinsic noise, arising from stochastic protein production, remains dominant at steady state, both sources (TFs, X and Y) significantly impact noise during the transient phase. This interplay between intrinsic and extrinsic noise explains the observed super-Poissonian behavior of protein levels in the transient regime. We note that the super-Poisson behavior, characterized by a variance exceeding the mean, is commonly observed in biological systems and has been previously reported in several studies [8,59,60].

The activation timing of genes significantly influences their communication and coordination. Symmetric activation (θ= 1) promotes information exchange between Z_1_ and Z_2_, leading to synchronized expression patterns. Conversely, asymmetric activation (θ≠ 1) disrupts this communication, resulting in less coordinated behavior. These effects are evident both at the instantaneous time point and across different time scales. The coefficient, θ, acts as a regulatory tool for tuning co-expression patterns and noise levels. While extreme values of θ can disrupt inter-gene communication and network functionality, intermediate values facilitate desired expression patterns. The optimal range of θ likely lies within a moderate interval, avoiding the extremes.

The asymmetric gene expressions can be advantageous and introduce incoherence in gene co-expression due to differential binding affinity of the shared TFs. Genes with stronger promoter affinity (higher binding affinity for the TF(s)/signal(s)) can detect weak, fluctuating signals more readily, leading to an earlier response with lower noise. The low noise level ensures a precise response compared to the delayed and potentially noisier response of genes with lower affinity promoters. The emerging hierarchy in noise and thus the generated phenotypic diversity allows the motif to adapt to diverse fluctuating signals. In contrast, symmetric expression promotes a coordinated response, ensuring they act together. Similar principles of asymmetric and temporal ordering in gene expression have also been observed in single-input regulatory modules present in phosphate starvation response in yeast [61,62]. The present analysis also highlights the role of network interconnection within TOFFLs compared to studying isolated FFLs. The interconnected nature provides a richer and more versatile toolbox for fine-tuning gene expression, allowing for a wider range of responses compared to isolated regulatory modules.

Our findings highlight the key role of asymmetric gene expression patterns, regulated by differential promoter affinity, in generating different phenotypes in the temporal limit. This mechanism allows for the production of diverse cellular responses, enhancing adaptability to fluctuating environments. This observation is consistent with a recent study demonstrating the importance of regulatory network architecture in conferring phenotypic plasticity — an organism’s capacity to exhibit different characteristics under diverse environmental conditions, in the quorum sensing system in *P. aeruginosa* [37]. The quorum sensing network employs a multi-output incoherent feed-forward loop and exemplifies how such networks can partition cellular responses while maintaining robustness to fluctuations in key regulatory factors. While our present study focuses on the coherent variant of the multi-output feed-forward loop (MOFFL), both coherent and incoherent MOFFL architectures, despite their distinct regulatory logics, share the capacity to generate phenotypic diversity through noise-driven mechanisms.

Analysis of the basic recurring structure, TOFFL, within the MOFFL network, provides deeper insights into how DBA shapes the timing of gene regulations and coordinates the behavior of gene products within complex transcriptional regulatory circuits. Various cellular processes require specific gene expression patterns characterized by precise timing and coordinated activation/deactivation between genes. Our findings suggest how cells can strategically modulate the binding strengths of TFs to achieve these desired patterns. We decipher the relationship between noise and network architecture during both transient and steady state dynamics. By examining how DBA influences noise in the co-expression of genes targeted by shared TFs, we gain valuable insights into how noise shapes the dynamics of biological processes. Noise can be both beneficial and detrimental [15,24,39,40], and our findings could inform the development of novel strategies for gene expression regulation, allowing for greater precision in tuning gene expression patterns by targeting specific TF binding affinities.

## ACKNOWLEDGEMENTS AND FUNDING SOURCES

Fruitful discussions with Smarajit Polley, Alok Kumar Maity, and Sandeep Choubey are thankfully acknowledged. MN thanks Science and Engineering Research Board (SERB), India, for the National Post-Doctoral Fellowship [PDF/2022/001807].

## Appendix A: Calculation of steady state moments

Eqs. (1-4) can be solved at steady state to obtain the covariance matrix (**σ**) associated with system components by writing down the continuous Lyapunov equation in the small noise limit [12,52,57],

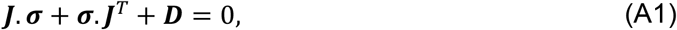

where *T* stands for the matrix transposition. The diagonal elements of the diffusion matrix, *D*, contain the noise strengths associated with the system variables and the off-diagonal elements are zero. In general, the matrix *D* is defined as ⟨ ξ_*i*_(*t*)ξ_*j*_(*t*^’^) ⟩ = [*f*_*i*_(⟨⋯ ⟩) + β_*i*_⟨*i*⟩]δ_*ij*_ δ(*t* − *t*^’^), evaluated at steady state. The elements of the Jacobian matrix, *J* are defined as,

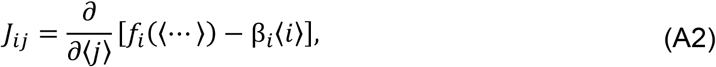

where ⟨⋯ ⟩ stands for the steady state ensemble average. We note that the Lyapunov equation (A1) is the result of standard linear noise approximation [49,52]. Solving Eq. (A1) gives the following variances and covariances,

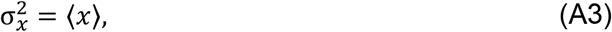

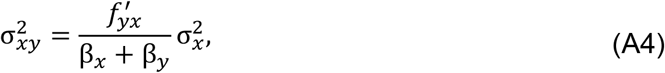

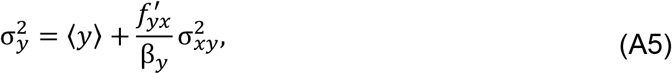

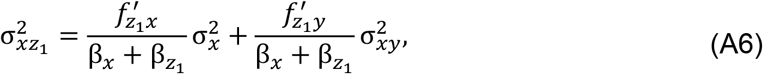

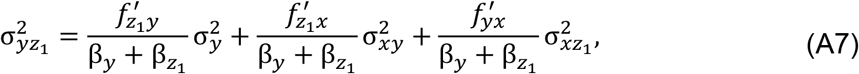

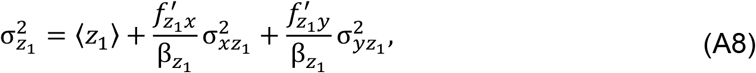

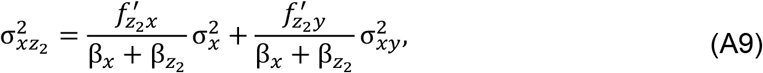

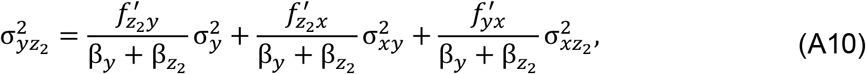

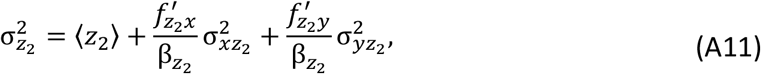

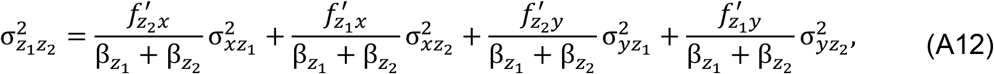

where, 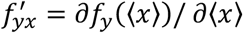, a partial derivative of the synthesis function evaluated at steady state. A similar notion of the definition applies to the other functional derivatives present in Eqs. (A6-A12). For the explicit form of the functions (evaluated at steady state), we refer to Table 1. We note that the presentation of steady state second moment calculations summarize the more detailed pedagogical derivations provided in our previous works [17,63,64].

We, now define the steady state FF 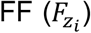 and 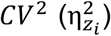 associated with the two output gene products, Z_1_ and Z_2_, as,

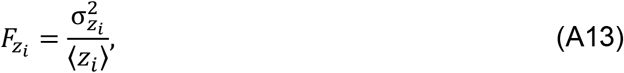

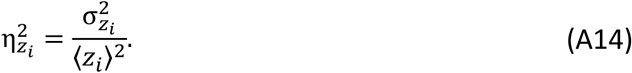

Substituting the expression of 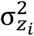 (Eqs. (A8) and (A11)) on Eq. (A13), we obtain the explicit forms of steady state intrinsic and extrinsic noise components [11,12,14,16],

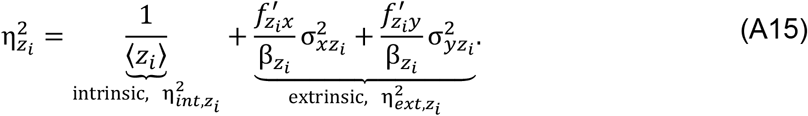

We rewrite the expression of FF (Eq. (A13)) in terms of *CV*^2^ (Eq. (A14)) and intrinsic-extrinsic parts as,

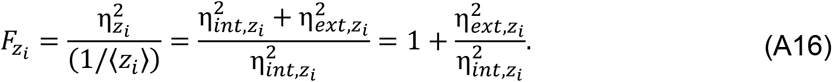

We note that this equation resembles Eq. (15) which correlates FF and noise decomposition in the transient limit.

We, now, define the steady state correlation coefficient and MI between Z_1_ and Z_2_ as,

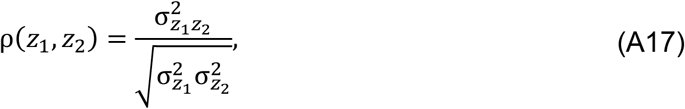

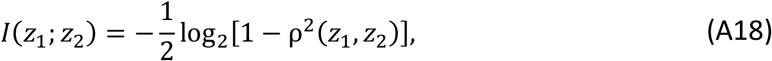

where we have used the Gaussian channel approximation to define the MI [58,65].

## Appendix B: Fano factor in terms of intrinsic and extrinsic noise at the transient limit

The temporal FF is defined as,

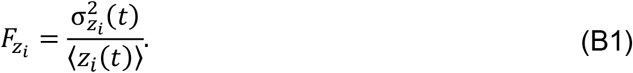

We, now, use Eq. (13) and (14) to rewrite FF in terms of intrinsic and extrinsic noise components as,

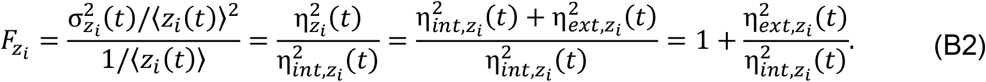

